# An important role for periplasmic storage in *Pseudomonas aeruginosa* copper homeostasis revealed by a combined experimental and computational modeling study

**DOI:** 10.1101/301002

**Authors:** Jignesh H. Parmar, Julia Quintana, David Ramírez, Reinhard Laubenbacher, José M. Argüello, Pedro Mendes

**Affiliations:** Center for Quantitative Medicine and Department of Cell Biology, University of Connecticut School of Medicine, 263 Farmington Av., Farmington, CT, 06030, USA; Department of Chemistry and Biochemistry, Worcester Polytechnic Institute, 100 Institute Road, Worcester, MA, 01609, USA; Jackson Laboratory for Genomic Medicine, 10 Discovery Dr., Farmington, CT, 06032, USA

**Author notes:** JHP and JQ should be considered joint first author. RL, JMA and PM should be considered joint senior author.

**Keywords:** *Pseudomonas aeruginosa*, copper, homeostasis, computational modeling, system identification

## Abstract

Biological systems require precise copper homeostasis enabling metallation of cuproproteins while preventing metal toxicity. In bacteria, sensing, transport and storage molecules act in coordination to fulfill these roles. However, there is not yet a kinetic schema explaining the system integration. Here, we report a model emerging from experimental and computational approaches that describes the dynamics of copper distribution in *Pseudomonas aeruginosa*. Based on copper uptake experiments, a minimal kinetic model describes well the copper distribution in the wild type bacteria but is unable to explain the behavior of the mutant strain lacking CopA1, a key Cu^+^ efflux ATPase. The model was expanded through an iterative hypothesis-driven approach, arriving to a mechanism that considers the induction of compartmental pools and the parallel function of CopA and Cus efflux systems. Model simulations support the presence of a periplasmic copper storage with a crucial role under dyshomeostasis conditions in *P. aeruginosa*. Importantly, the model predicts not only the interplay of periplasmic and cytoplasmic pools but also the existence of a threshold in the concentration of external copper beyond which cells lose their ability to control copper levels.

## Introduction

Copper is an essential trace element used as a cofactor in redox enzymes (Fraústo da Silva and Williams, 2001; Cobine *et al*., 2006; Tavares *et al*., 2006). However, copper in its “free” form is highly toxic and leads to mismetallation of proteins, disassembly of Fe-S centers, and generation of free radicals (Macomber and Imlay, 2009; Dupont *et al*., 2011). Therefore, organisms have evolved molecular systems targeting the metal to cuproproteins while limiting high intracellular copper levels. Understanding of bacterial mechanisms of copper homeostasis is relevant in light of their importance for pathogenic virulence. The antibacterial properties of copper and role in innate immunity are well established (Hodgkinson and Petris, 2012; Ladomersky and Petris, 2015; Djoko *et al*., 2015). Studies on bacterial response to copper stress have revealed various essential components such as sensing transcriptional regulators, membrane transporters and chaperones that control Cu^+^ levels and localization (Rensing and Grass, 2003; Osman and Cavet, 2008; Argüello *et al*., 2013). Deletion of some of these proteins leads to a consequent copper dyshomeostasis. In addition proteins that bind copper with high capacity and might act as storage pools have been proposed recently (Vita *et al*., 2016). Nevertheless, there is still a lack of a general model to describe and predict the accumulation and distribution of copper in Gram-negative bacteria in kinetics terms.

Several proteins that play a role in copper sensing, transfer, and distribution between compartments have been identified in the periplasm, cytoplasm, inner, and outer membranes of *Pseudomonas aeruginosa* (Yoneyama and Nakae, 1996; Teitzel *et al*., 2006; Caille *et al*., 2007; Frangipani and Haas, 2009; Thaden *et al*., 2010; González-Guerrero *et al*., 2010; Elsen *et al*., 2011; Quintana *et al*., 2017). Extracellular Cu^2+^ likely enters the periplasm through OprC, an outer membrane porin (Yoneyama and Nakae, 1996). While OprC binds Cu^2+^, the specificity of transport as well as the putative presence of external reductases generating Cu^+^ has not been established. The inner membrane CcoA transports into the cytoplasm the periplasmic copper destined to metalate cytochrome c oxidase (Ekici *et al*., 2012), while novel members of the Major Facilitator Superfamily might also enable copper influx through this membrane (Quintana *et al*., 2017). Copper is likely supplied to cytoplasmic cuproproteins and other putative copper binging pools via cytoplasmic chaperones, as has been proposed for the role of chaperones in delivering the metal to inner membrane Cu+ exporters (P1B-ATPases) and transcriptional sensors (Cobine *et al*., 1999; González-Guerrero *et al*., 2009). Two cytoplasmic Cu^+^ chaperones, CopZ1 and CopZ2, have been identified in *P. aeruginosa* although their singular roles have not been established (Quintana *et al*., 2017). Two Cu^+^-ATPases, CopA1 and CopA2, mediate cytoplasmic copper export in these bacteria. CopA2 provides copper for cytochrome c oxidase, while CopA1 functions as a primary exporter to the periplasm, guarding against excess buildup of cytoplasmic Cu^+^ (González-Guerrero *et al*., 2010). Cellular copper might also be exported by the *P. aeruginosa* Cus system, CusA, CusB and CusC, spanning from the inner to the outer membrane (Thaden *et al*., 2010; Kim *et al*., 2011; Quintana *et al*., 2017). This efflux system has been proposed to transport copper from the cytoplasmic and periplasmic copper pools to the extracellular space in *E. coli* (Kim *et al*., 2010; Long *et al*., 2010; Kim *et al*., 2011). However, *P. aeruginosa* does not have a periplasmic chaperone CusF that might deliver copper to CusB (Kim *et al*., 2010; Quintana *et al*., 2017), suggesting that cytoplasmic Cu^+^ might be the substrate of this Cus system. Regarding periplasmic copper, this is likely exported through the outer membrane protein PcoB (Cha and Cooksey, 1991; Lee *et al*., 2002). Expression of these transporters and chaperones, as well as other proteins of unknown functions, is controlled by a two component periplasmic copper sensor (CopS/CopR) and the cytoplasmic sensor CueR (Thaden *et al*., 2010; Quintana *et al*., 2017).

Using genome-wide transcriptomic analysis, we have proposed an integrated copper homeostatic network (Quintana *et al*., 2017). This considers the requirement of specific influx and efflux systems and subcellular metal chaperones; as well as transcriptional regulation of the various functional elements in each compartment. However, these observations do not explain how the interactions of the metal distribution components function in a coordinated manner to timely respond to environmental changes. Mathematical models are increasingly used to understand complex biological systems and test different mechanisms that can explain the behavior at system level. Previously developed mathematical models of copper homeostasis in archaea and Gram-positive bacteria (Pécou *et al*., 2006; Pang *et al*., 2013) have not addressed the dynamics of Cu^+^ movements via *bona fide* transporters and chaperones nor the distribution across periplasmic and cytoplasmic pools. We hypothesized that mathematical analysis of copper uptake kinetics might provide evidence on the necessary elements (influx and efflux transporters and compartmental pools) that explain the organization of the reported homeostatic network. Here we present a predictive mathematical model, consistent with experimental observations, that advances our understanding of the mechanisms of copper homeostasis in *P. aeruginosa*.

## Results

### Modeling copper uptake kinetics

Our goal was to develop a mathematical model that can describe the copper uptake kinetics at various concentrations of extracellular metal, as well as the distribution in the two cellular compartments, periplasm and cytoplasm. We expected that simulations can predict the copper homeostasis not only for the wild type strain but also under conditions of dyshomeostasis; i.e. when an element of the system is knocked out via mutagenesis. We have previously shown that *P. aeruginosa* viability and growth rate are not compromised by external copper concentrations up to 0.5 mM Cu^2+^, while incubation at higher levels leads to eventual cell death (Quintana *et al*., 2017). In the same work, copper uptake kinetics at this “threshold” concentration was also studied. These data served for an initial assessment of the proposed model (Figure 1A). Nonetheless, towards enhancing the scope of our approach, the compartmental distribution was estimated at pre-steady (1 min, 10 min) and steady state (30 min) (Fig. 1B). We also analyzed Cu^2+^ uptake kinetics of wild type *P. aeruginosa* in the presence of 2 mM Cu^2+^ (Fig. 1C) and 4 mM Cu^2+^ (Fig. 1D) in the culture medium. Cell viability started to decrease in cells exposed to 2 mM CuSO_4_ after 1 h treatment, whereas toxicity is evident as soon as 10 min after the addition of 4 mM CuSO_4_ (data not shown). Cells exposed to 2 mM Cu2+ showed a modest but noticeable increase in Cu levels beyond the initial fast metal uptake. Alternatively, exposure to 4 mM Cu2+ lead to a marked steady increase of intracellular metal levels over the course of the experiment. Then, it is apparent the presence of a “switch” from tolerance to toxicity at around 1 attomol of copper per cell (Fig. 1C), as higher steady state levels lead to cell death (Quintana et al., 2017).

**Figure 1.**
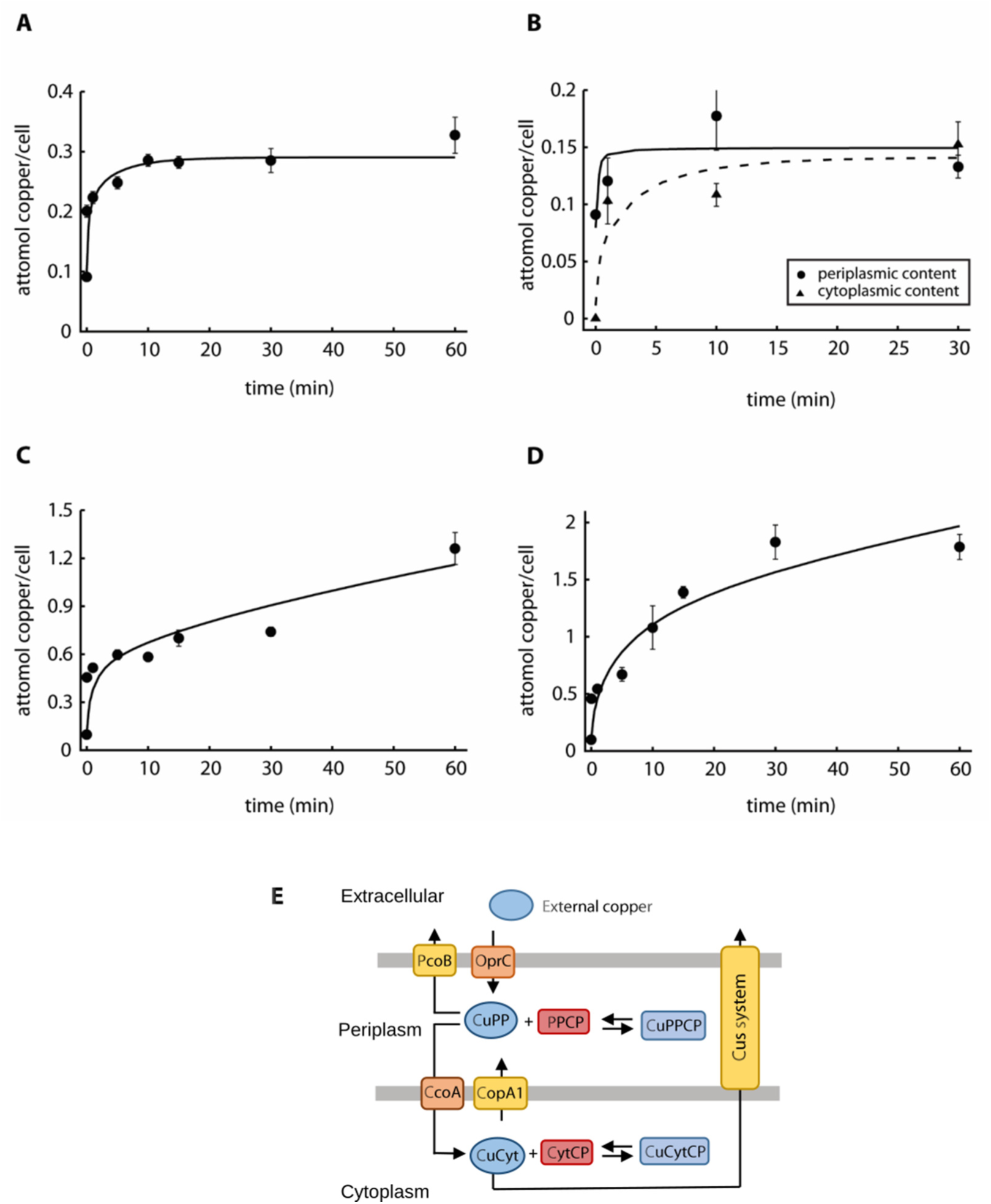
Experimental data and best fit by Model I. A) Copper uptake kinetics in the presence of 0. 5 mM CuSO_4_ (Quintana et al. 2017). B) Compartmental distribution of copper levels 1, 10 and 30 min after the addition of 0.5 mM CuSO_4_, solid line is for periplasmic and dashed line is for cytoplasmic fraction. C, D) Copper uptake kinetics in the presence of 2 mM and 4 mM CuSO_4_. Solid lines represent the best fit to the data using Model I (parameter values in Table 1). Experimental data are the mean ± standard error of 3 independent replicates. E) Parsimonious model of copper uptake kinetics in P. aeruginosa PAO1 (Model I). CuPP, periplasmic copper bound to chaperones, and CuCyt, cytoplasmic copper bound to chaperones. PPCP, periplasmic and outer membrane proteins; CytCP cytoplasmic and inner membrane proteins. Equilibrium copper binding is represented by arrows pointing both ways. Single arrows represent the direction of transport.

We initiated the modeling of the uptake kinetics using a parsimonious Model I (Fig. 1E). A fixed total cell volume of 10^−15^ L is adopted in the model, composed of a periplasmic compartment (20% of the total volume) and a cytoplasmic compartment (80% of the total volume) (Kumar and Imlay, 2013). For modeling purposes, based on the virtual absence of free (unbound) copper in the cell (Changela *et al*., 2003; Smirnova *et al*., 2012), we assumed all copper to be bound to proteins (copper chaperones and cuproproteins). We use the term copper chaperones to refer to those cuproproteins that bind copper as a means to hand it off/on transporters. We term cuproproteins those proteins that have copper bound for a long time scale and do not make it available to transport until it dissociates and binds to chaperone. Molecular elements of the model could be predicted based in our earlier transcriptional analysis; i.e. chaperones CopZ1 and CopZ2, transporters CcoA and CopA1, etc. However, given the uncertainties on the identity of periplasmic chaperones, as well as on the roles of both cytoplasmic chaperones (CopZ1 and CopZ2), the species *CuPP* and *CuCyt* represent Cu^+^ bound to periplasmic and cytoplasmic chaperone molecules respectively (Fig 1E). To reduce the model complexity all cuproproteins were grouped into one pool per compartment; *PPCP* represents all periplasmic and outer membrane cuproproteins, and *CytCP* represents all cytoplasmic and inner membrane cuproproteins. It was considered that Cu^+^ binding to these proteins follows mass action kinetics. For simplicity, the model does not explicitly consider the apo forms of the chaperones; this is, their concentration is not limiting. The Cu^+^ binding and dissociation with cuproproteins (*PPCP* and *CytCP*) is also represented by mass action, however in this case we represent explicitly the apo version of the proteins. Our experimental results suggested that before cells are exposed to Cu2+ (t = 0), most of intracellular copper exists in the periplasm (Fig. 1B). However, in order to undertake experimental constraints, we allowed the initial concentration of cytoplasmic copper to vary between 0 to 30% of the total during parameter estimation. Then, the initial concentrations of copper bound to cuproproteins (CuPPCP and CuCytCP) were obtained through parameter estimation while the concentrations of copper bound to chaperones (CuCyt and CuPP) were obtained by subtracting those estimated values from the total compartmental copper. The initial concentrations of PPCP and CytCP were also parameters estimated to fit the data. We assumed that extracellular Cu^2+^ *(CuExt)* enters the periplasm *(CuPP)* through the influx protein (OprC) and exported via an efflux protein (PcoB). Periplasmic and cytoplasmic Cu^+^ exchange is through dedicated influx and efflux transporters (CcoA and CopA). In addition, we included in the model the export of cytoplasmic Cu^+^ into the medium via the Cus-system present in *P. aeruginosa*. Considering the different transmembrane systems, their transport rates were defined by Michaelis-Menten equations. The resulting differential equations for transport and equilibrium copper exchange were integrated in a full model (see Supplemental Information) that yielded parameter values (Table 1) that fit well the experimental Cu^+^ uptake curves in a range of external Cu^2+^ concentrations (Fig. 1A, 1C, 1D), as well as the compartmental distribution (Fig. 1B).

**Table 1.** - Best fit parameter values for Models I, II, and III.

| Parameter | Role | Best fit value |  |  | Units |
| --- | --- | --- | --- | --- | --- |
|  |  | Model I | Model II | Model III |  |
| vOprC | Limiting rate of copper import from medium to periplasm | 7.83 | 2.44 | 6.84 | mmol/min |
| KmOprC | Michaelis constant for copper import from medium to periplasm | 0.546 | 0.328 | 1.66 | mM |
| vPcoB | Limiting rate of copper export from periplasm to medium | 7.30 | 2.20 | 3.68 | mmol/min |
| KmPcoB | Michaelis constant for copper export from periplasm to medium | 0.33 | 0.232 | 0.390 | mM |
| vCcoA | Limiting rate for copper import from periplasm to cytoplasm | 2.72 | $1.19 \times 10^6$ | 4.43 | mmol/min |
| KmCcoA | Michaelis constant for copper import from periplasm to cytoplasm | 22.1 | 36.7 | 9.50 | $\mu\text{M}$ |
| vCopA1 | Limiting rate for copper export from cytoplasm to periplasm | 0.665 | 0.127 | 0.713 | mmol/min |
| KmCopA1 | Michaelis constant for copper export from cytoplasm to periplasm | 6.00 | 5.42 | 0.476 | $\mu\text{M}$ |
| kasCuPPCP | On constant for copper to periplasmic cuproproteins | 595.3 | 283 | $6.98 \times 10^4$ | 1/M min |
| kdsCuPPCP | Off constant for copper from periplasmic cuproproteins | $2.24 \times 10^{-6}$ | $1.31 \times 10^{-6}$ | $1.26 \times 10^{-6}$ | 1/min |
| kasCuCytCP | On constant for copper to cytoplasmic cuproproteins | $4.19 \times 10^8$ | $8.33 \times 10^8$ | $4.70 \times 10^8$ | 1/M min |
| kdsCuCytCP | Off constant for copper from cytoplasmic cuproproteins | 97.9 | 98.4 | 98.6 | 1/min |
| vCusSystem | Limiting rate for copper export from cytoplasm to medium | 1.02 | 1.23 | 0.390 | mmol/min |
| KmCusSystem | Michaelis constant for copper export from cytoplasm to medium | 33.4 | 0.337 | 38.9 | $\mu\text{M}$ |
| kdPPCP | First order rate constant for periplasmic cuproproteins degradation (turnover) | NA | NA | 0.005 | 1/min |
| ksPPCP | Limiting rate for synthesis of periplasmic cuproproteins | NA | NA | 14.4 | $\mu\text{mol/min}$ |
| KaCuCytPPCP | Activation constant by copper towards synthesis of periplasmic cuproproteins | NA | NA | 0.0380 | $\mu\text{M}$ |
| vCopA2 | Limiting rate for copper export from cytoplasm to periplasm through CopA2 | NA | $5.46 \times 10^5$ | NA | mmol/min |
| kmCopA2 | Michaelis constant for copper export from cytoplasm to periplasm through CopA2 | NA | 0.139 | NA | $\mu\text{M}$ |

The resulting kinetic parameter values (Table 1) are within expected transport rates and equilibrium affinity constants (González-Guerrero *et al*., 2008; González-Guerrero *et al*., 2010; Xiao *et al*., 2011; Argüello *et al*., 2012; Quintana *et al*., 2017). Importantly, the model can describe the copper distribution between the periplasmic and cytoplasmic compartments (Fig. 1E) and marks the parallel transport by the CopA and Cus-system. In this direction, exclusion of the Cus-system prevented fitting of the uptake curves.

### Effects of dyshomeostasis on copper uptake kinetics

Previous results in our laboratory showed that a *P. aeruginosa* mutant strain lacking CopA1 (*ΔcopA1*) has increased intracellular Cu^+^ levels that result in an increase sensitivity to the metal (González-Guerrero *et al*., 2010). To aid the mathematical modeling of copper distribution, we studied the kinetics of copper accumulation in *ΔcopA1* after exposure to 0.5 mM CuSO_4_ as well as the metal compartmental distribution (Fig. 2). We previously observed that *P. aeruginosa*, as well as mutant strains lacking periplasmic (*ΔcopR*) and cytoplasmic (*ΔcueR*) Cu^+^ sensors, growth at normal rates at the sub-lethal concentrations of 0.5 mM external copper (Quintana *et al*., 2017). A similar behavior was observed in the *ΔcopA1* strain and this allowed the study of copper levels per cell. Determination of the copper uptake kinetics showed metal levels rose faster in the mutant strain than in the wild type bacteria reaching steady state after 60 min (Fig. 2A; compare to Fig. 1A). As previously shown, mutant complementation with the *copA1* gene reestablished the wild type phenotype (González-Guerrero *et al*., 2010). Interestingly, the compartmental response of the mutant strain was distinct from that of the wild type and relatively faster overload of the periplasm compared to cytoplasm was observed (Fig. 2B). Our minimal Model I was incompatible with these experimental data as it predicted an increase of the cytoplasmic pool. This is, it was not possible to obtain good global fits against the WT and *ΔcopA1* data with the minimal model. We then used the parameter values obtained just using WT data to simulate the *ΔcopA1* mutant by setting the *V_COPA1_* parameter to zero; the results predicted that the cytoplasm would accumulate very high copper concentration while the periplasmic copper would decrease below its initial concentration. This contradicts the experimental results suggesting the presence of additional homeostatic components.

**Figure 2.**
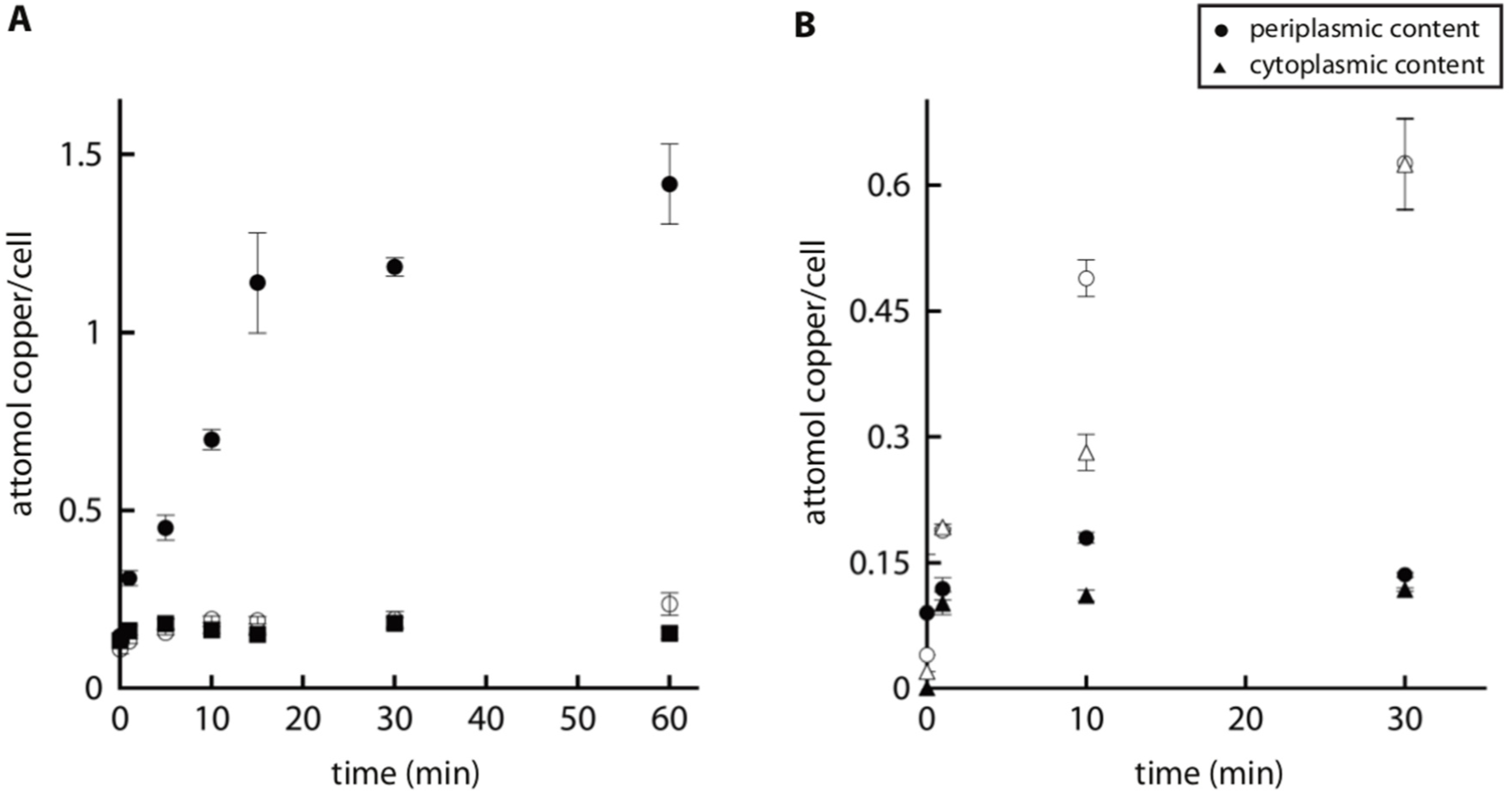
Effect of dyshomeostasis on copper uptake kinetics in *P. aeruginosa* PAO1. Copper uptake kinetics in the insertional mutant ΔcopA1 (black circles) and the corresponding complemented strain (black squares) after addition of 0.5 mM CuSO_4_. Empty circles correspond to WT. B) Compartmental distribution of copper levels in the insertional mutant ΔcopA1 (white shape) and WT (black shape) 1, 10 and 30 min after the addition of 0.5 mM CuSO_4_. Experimental data are the mean ± standard deviation of 3 independent replicates.

Looking for an alternative model, we considered that *P. aeruginosa* has two Cu^+^-ATPases (CopA1 and CopA2) responsible for cytoplasmic Cu^+^ efflux (González-Guerrero *et al*., 2010). The rationale behind this hypothesis was that if CopA2 could transport copper at significant higher rate from cytoplasm to periplasm in the absence of CopA1 then the periplasm would accumulate more copper compared to the cytoplasm. A new Model II was therefore created to include two cytoplasm to periplasm efflux systems (Fig. 3A). Model II was able to fit the behavior of the wild type strain (Fig. 3B, 3C, 3C, 3D). However, Model II also failed to describe the behavior of the Δ*copA1* mutant strain. Although only *copA1* is apparently regulated by copper (González-Guerrero *et al*., 2010; Quintana *et al*., 2017), a variation of Model II was tested where the *V_CopA2_* parameter was allowed to increase with the cytoplasmic copper concentration, as if transcriptionally upregulated. However, this modification still did not lead to the satisfactory fitting of the data.

**Figure 3.**
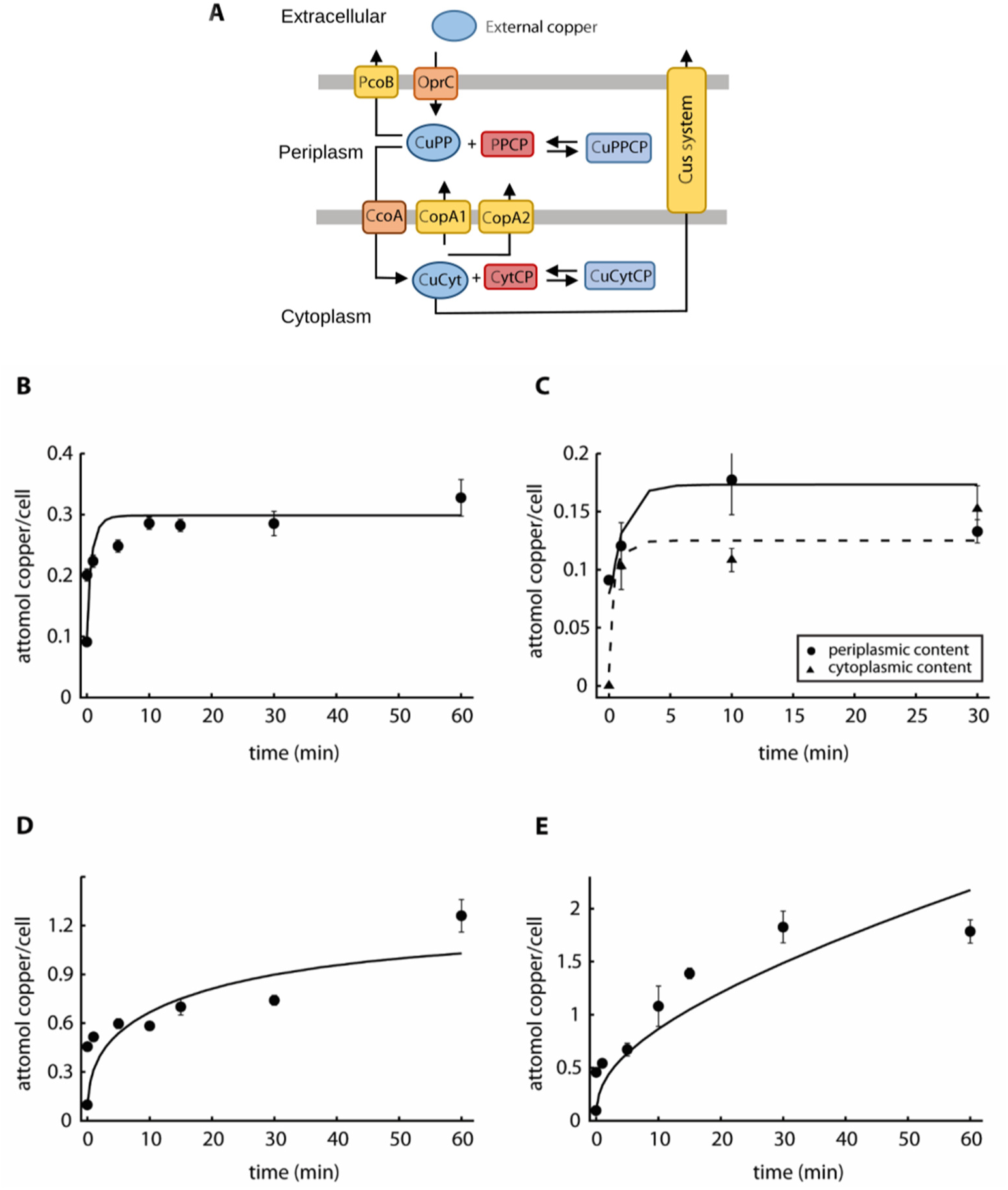
Experimental data and best fit by Model II. A) Modified model, including CopA2, of copper uptake kinetics in P. aeruginosa WT PAO1 (Model II). CuPP, periplasmic copper bound to chaperones, and CuCyt, cytoplasmic copper bound to chaperones. PPCP, periplasmic and outer membrane proteins; CytCP cytoplasmic and inner membrane proteins. Equilibrium copper binding is represented by arrows pointing both ways. Single arrows represent the direction of transport. B) Copper uptake kinetics in the presence of 0.5 mM copper (Quintana et al. 2017). C) Compartmental distribution of copper levels 1, 10 and 30 min after the addition of 0.5 mM CuSO_4_, solid line is for periplasmic and dashed line is for cytoplasmic fraction. D, E) Copper uptake kinetics in the presence of 2 mM and 4 mM CuSO_4_. Solid lines represent the best fit to the data using Model II (parameter values in Table 1). Experimental data are the mean ± standard deviation of 3 independent replicates.

**Figure 4.**
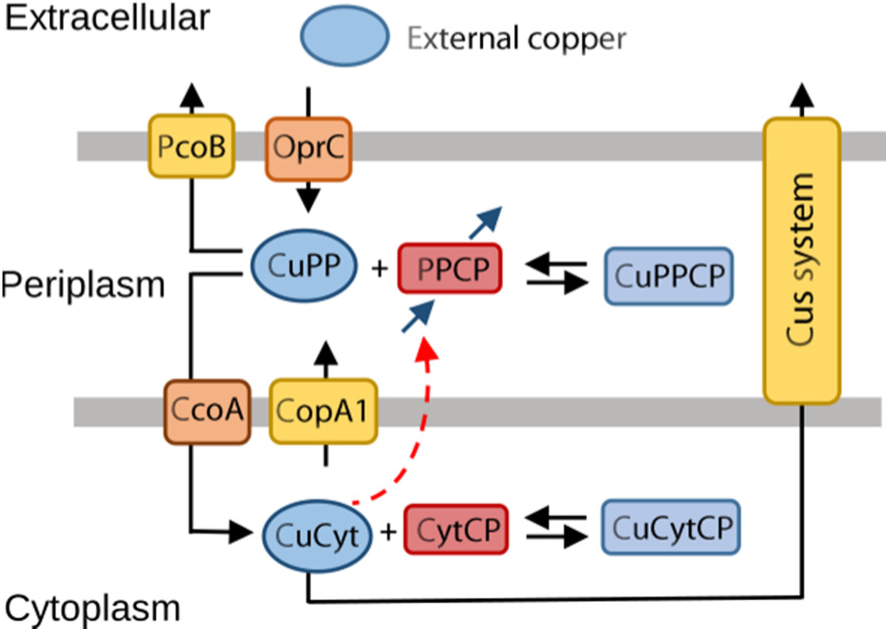
Model explaining the role of periplasmic copper pool in *P. aeruginosa* PAO1 (Model III). CuPP, periplasmic copper bound to chaperones, and CuCyt, cytoplasmic copper bound to chaperones. PPCP, periplasmic and outer membrane proteins; CytCP cytoplasmic and inner membrane proteins. Black arrows represent transport and binding, blue arrows represent synthesis and degradation of PPCP and red dashed arrow represents induction of PPCP synthesis by CuCyt. Note that synthesis of PPCP happens in the cytoplasm and the protein is appropriately transported to the periplasm, but for clarity the synthesis arrow is included in the periplasm in this diagram.

An alternative mechanism that could explain the higher periplasmic copper levels in the ΔcopA1 strain could be an increased expression of the periplasmic cuproproteins. Supporting this hypothesis, we have observed high levels of periplasmic cuproproteins in the ΔcopA1 albeit determinations were performed in the absence of Cu2+ stress (Raimunda *et al*., 2012). Following this line of thinking, gene expression would be increased in the cytoplasm after sensing of a high copper level in the periplasm by CopS/R, to ultimately induce higher synthesis of periplasmic cuproproteins. Testing this hypothesis, we created Model III where the periplasmic pool of cuproproteins *(PPCP)* is induced by the cytoplasmic copper *(CytCP)* (Fig. 4). Here, we also represent the synthesis and degradation of *PPCP;* the degradation rate of *PPCP* is assumed to be a first order process, while the synthesis rate is assumed to be a sigmoidal function of *[CuCyt]*:

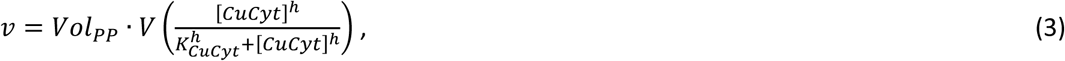

where *V* is a basal rate of synthesis, *h* is a Hill coefficient, *K_CuCyt_* is an affinity constant for the activation of the synthesis rate, and *Vol_PP_* is the volume of the periplasm. We then proceeded to fit this model against both the wild type and the *ΔcopA1* strains data. Interestingly, Model III was able to simultaneously reproduce the response from both *ΔcopA1* and wild type strains (Fig. 5A, 5C, 5D, 5E), including the compartmental distribution (Fig. 5B, 5F). Moreover, the model accurately predicted the steady-state conditions for WT after 5-10 min (Fig. 5A, 5C, 45D) and the increase in copper levels in *ΔcopA1* even 60 min after exposure (Fig. 5E).

**Figure 5.**
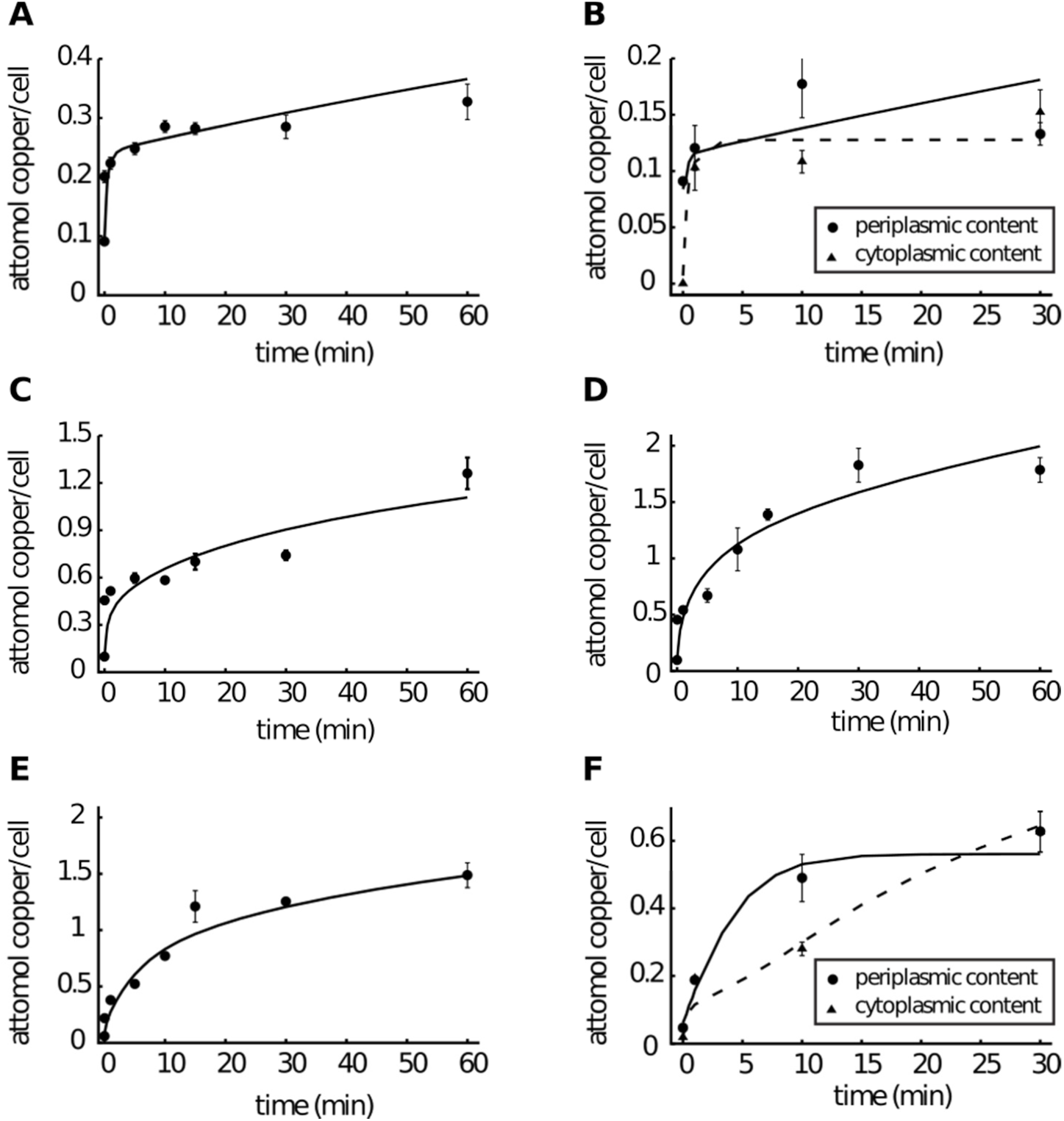
Experimental data and best fit by Model III. A) Copper uptake kinetics in the presence of 0.5 mM CuSO_4_ (Quintana et al. 2017). B) Compartmental distribution of copper levels 1, 10 and 30 min after the addition of 0.5 mM CuSO_4_, solid line is for periplasmic and dashed line is for cytoplasmic fraction. C, D) Copper uptake kinetics in the presence of 2 mM and 4 mM CuSO_4_. E) Copper uptake kinetics of the insertional mutant ΔcopA1 in the presence of 0.5 mM copper. F) Compartmental distribution of copper levels in the insertional mutant ΔcopA1 1, 10 and 30 min after the addition of 0.5 mM CuSO_4_, solid line is for periplasmic and dashed line is for cytoplasmic fraction. Solid lines represent the best fit to the data using Model III (parameter values in Table 1). Experimental data are the mean ± standard deviation of 3 independent replicates.

For each of the models we characterized parameter identifiability using likelihood profiles (Raue *et al*., 2009). As depicted in Figures S1-S2, most parameters of the Models I and II are not identifiable (those that have flat curves), the only exception being *KmOprC* and some of the initial concentrations. However, in the case of the Model III (Figure S3), most parameters are identifiable within 90% confidence intervals. The dissociation rate constants *kdsCuPPCP* and *kdsCuCytCP* are not identifiable for all three models because they can be counterbalanced by the respective association constants (to determine these parameters well would require binding studies with purified proteins). The other non-identifiable parameter, common to all three models, is *KmPcoB* which suggests that the export of copper from periplasm through the outer membrane may not be essential to explain the experimental observations. Overall, the analysis indicated that the inclusion of *ΔcopA1* mutant data in Model III is what constrained the model parameters to become identifiable. This provides further confirmation that Model III is the best of the three in describing copper transport and homeostasis, and suggests a prominent role of the periplasm in protecting from higher environmental copper.

### Model Predictions

One of the advantages of computational modeling is the capability to simulate experiments where predictions can be tested. Toward this goal, we used Model III to simulate the *P. aeruginosa* response of external copper concentrations higher than 4 mM (Fig. 6). Model III suggests that, in these conditions, intracellular copper accumulation continuously increases without attaining a steady state (Fig. 6A). This is *P. aeruginosa* loses its homeostatic control above a threshold of 2-4 mM Cu^2+^ leading to intracellular metal levels above 1 attomol/cell. In the case of the *ΔcopA1* strain, this extracellular Cu^2+^ threshold is even lower since the internal copper increases at faster rate (Fig. 6B). This is in agreement with the experimental observation that the *ΔcopA1* strain is much more sensitive than wild type *P. aeruginosa* to high levels of extracellular Cu^2+^ (González-Guerrero *et al*., 2010). Moreover, Model III predicts that the periplasmic copper pool becomes larger than the cytoplasmic at 1 mM external Cu^2+^. However, at highly toxic external copper conditions, i.e. 4 mM, cytoplasmic copper increases dramatically compared to the periplasmic concentration (Fig. 6B). On the other hand, the *ΔcopA1* strain accumulates high copper in the cytoplasm at both 1 mM and 4 mM external copper (Fig. 6C). Thus, the model predicts that *P. aeruginosa* respond to increased copper initially inducing periplasmic storage. However, beyond a certain concentration of external copper, tolerance mechanisms are saturated and copper is re-distributed, i.e. cytoplasmic storage mechanisms are induced. This threshold is lower when copper homeostasis is impaired, as it occurs in *ΔcopA1*.

**Figure 6.**
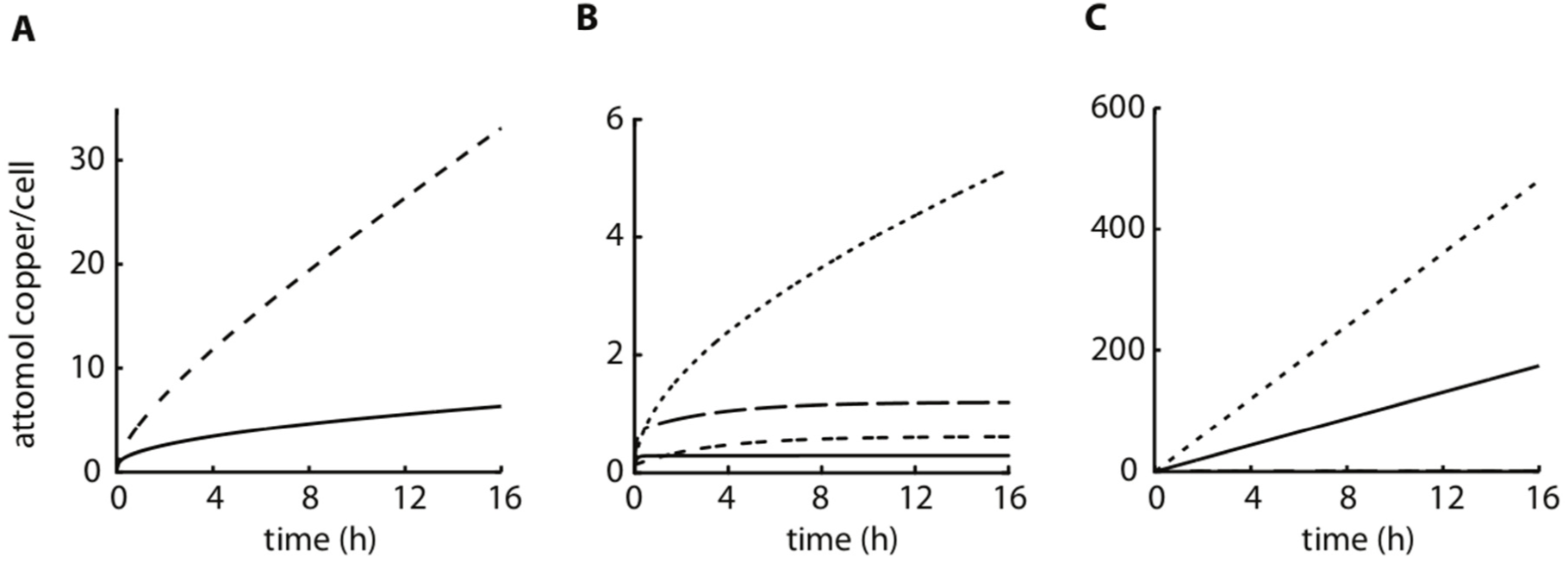
Predicting the effects of copper toxicity using Model III. A) Computational prediction of total cellular copper accumulation in *P. aeruginosa* WT strain for 4 mM (solid line) and 6 mM (dashed line) CuSO_4_. B) Computational prediction of copper levels in the *P. aeruginosa* WT strain in the presence of 1 mM and 4 mM CuSO_4_ in the periplasm (dashed line for 1 mM and dashed-dotted line for 4 mM) and for the cytoplasm (solid line for 1 mM and dotted line for 4 mM). C) Computational prediction of copper levels in the *P. aeruginosa* Δ*copA1* strain in the presence of 1 mM and 4 mM CuSO_4_ in the periplasm (dashed line for 1 mM and dashed-dotted line for 4 mM) and for the cytoplasm (solid line for 1 mM and dotted line for 4 mM; note that the periplasm concentrations are very small).

## Discussion

To shed light on the mechanisms of copper homeostasis in *P. aeruginosa*, we have previously established protein components (transporters, chaperones, transcriptional regulators) that enable metal uptake, distribution and efflux (Quintana *et al*., 2017). Along this line, we reported the regulons of periplasmic and cytoplasmic metal sensors that control the copper levels in each cellular compartment. However, this system description did not provide a quantitative explanation of the dynamic interplay among these homeostatic elements. Here, we present a mathematical model that using a single set of parameters is able to replicate the observed copper membrane fluxes and compartmental distribution in *P. aeruginosa*. Moreover, this model is able to describe the system under conditions of dyshomeostasis by exposure to high copper or deletion of a key protein.

Our strategy employed a bottom-up modeling approach starting with a minimal Model I. This considered all the transmembrane transporters identified in *P. aeruginosa* and pooled copper chaperones and cuproproteins into single compartmental pools. Departing from a parsimonious Model I, mechanistic complexity was expanded and tested against experimental observations. The proposal of mechanistic hypotheses at each step led to a minimal model that is consistent with the observations. Moreover, this iterative approach reduced the possibility of over-parameterization and data overfitting.

As the experimental basis for system simulations, we analyzed the accumulation and intracellular distribution of copper in *P. aeruginosa* exposed to increasing external metal concentrations, both in wild type and Δ*copA1* strains. Three kinetic models representing different mechanisms were tested in their capability to reproduce copper uptake data. Two of these models were unable to simultaneously explain the experimental observations from the two strains, indicating that the hypothetical systems were not consistent with the cellular response. Thus, we rejected the minimal Model I (Fig 1E) unable to simulate the copper compartmental distribution in the *ΔcopA1* strain; as well as the Model II where CopA2 would not undertake the function of CopA1 when the latter is deleted (Fig. 3A). Model III, which includes an induction of periplasmic cuproproteins by cytoplasmic copper is able to fit, with a single set of parameter values, all of our experimental data. Indeed we found out that not only is Model III better at fitting the experimental observations, but it also has better parameter identifiability. This may seem surprising since Model III is more complex than the two other models. In this case the extra complexity added in Model III was important in constraining the parameter values. This also gives validity to the approach adopted here: starting with simple models and gradually adding complexity until a good fit and identifiability is obtained.

The final Model III supported hypothetical and experimental observations. The model was consistent with the need of specific metal influx transporters in the outer and inner membranes. The outer membrane porin OprC, down regulated in the presence of high copper, appears as the logical candidate for influx transport across this membrane (Quintana *et al*., 2017). We assumed that CcoA mediates transport across the inner membrane (Ekici *et al*., 2012). However, required for metallation of cytochrome c oxidase, CcoA is not down regulated in conditions of metal stress and it is likely that a member of the Major Facilitator Superfamily encoded by PA5030 is a major contributor to copper influx (Quintana *et al*., 2017). Regarding copper efflux, Model III supports two significant characteristics of the system. First, it is congruent with the previously reported roles of CopA1 maintaining cytoplasmic copper levels and CopA2 supplying the metal for metallation of cytochrome c oxidase (González-Guerrero *et al*., 2010). Further studies are required to understand the integration of CcoA, CopA2 and perhaps specific chaperones in the metallation of respiration complexes. Second, the model points to the parallel function of a CopA1-PcoB tandem and the Cus-system to control the cell copper levels. While it appears to be a larger contribution of CopA1 relative to Cus (compare estimated V_max_), the transport via Cus appears essential to explain the experimental observations.

As expected, the model describes well the cellular metal accumulation which could be simulated assuming the absence of free copper. Importantly, it revealed an unexpected important early role of the periplasmic compartment in controlling copper levels. This is mediated by the upregulation of the periplasmic pool of cuproproteins (*PPCP)*. The proposed model formally assumes that a cytoplasmic cuproprotein is responsible for the induction of the periplasmic cuproprotein pool. If the present hypothesis is correct, deletion of this protein (likely CopR) should result in phenotypes severely affected in copper homeostasis.

We have previously reported that azurin (PA4922) is the major cupropotein in the periplasm of *P. aeruginosa* (Raimunda *et al*., 2013). Moreover, the expression of *azu* is increased in the Δ*copA1 and* Δ*copA2* stains compared to wild type strain. However, rather than having a metallochaperone role azurin has been reported to serve as an electron transfer carrier redox protein (Gray and Winkler, 1996), then other periplasmic proteins regulated by the CopS/R system might serve as copper sinks. In this direction, we observed the upregulation of a cupredoxin-like protein (PA2807), also with potential redox activity and multicopper oxidase PcoA (Djoko *et al*., 2008). The upregulation of several periplasmic redox enzymes not only supports the proposed role of oxidases producing a “less-toxic” Cu(II) (Grass and Rensing, 2001), but provide a molecular framework for the early accumulation of periplasmic Cu upon copper stress.

It is also notable that Model III does not include changes in the level of the cytoplasmic pool of cuproproteins. However this is because of the induction of periplasmic cuproproteins forms a negative feedback which stabilizes the cytoplasmic copper pool. Indeed the cytoplasm is an important compartment for copper sensing, distribution, and storage in bacteria (Padilla-Benavides *et al*., 2013; Philips *et al*., 2015; Vita *et al*., 2015). We have been unable to identify a cytoplasmic storage protein in *P. aeruginosa*, since no change in expression of likely candidates (metallothionein, Csp3, or glutathione biosynthesis genes) is observed during high-copper stress (Quintana *et al*., 2017).

In summary, we present a mathematical model that explains the dynamic function of influx and efflux systems in coordination with periplasmic and cytoplasmic metalloprotein copper pools toward achieving copper homeostasis. Importantly, the established parameters can simulate conditions of dyshomeostasis by exposure to high metal levels and the removal of system components.

## Experimental procedures

### Bacterial strains and growth conditions

*P. aeruginosa* PAO1 wild type and PW7626 (*PA3920/copA1* insertional mutant, *ΔcopA1)* strains were obtained from the Comprehensive *P. aeruginosa* Transposon Mutant Library at the University of Washington Genome Center (Jacobs *et al*., 2003). The *ΔcopA1* strain was complemented with the copA1 gene under control of its own promoter region as described (González-Guerrero *et al*., 2010). Cells were grown at 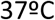 in Luria-Bertani (LB) medium, supplemented with irgasan (25 μg/ml), tetracycline (60 μg/ml), or kanamycin (35-50 μg/ml) as required.

### Copper uptake assays

Copper uptake was measured using cells in mid-log phase in antibiotic-free LB medium. Upon addition of 0.5, 2, or 4 mM CuSO4, aliquots were taken at indicated times for cell counting and copper levels determinations. Metal was measured in cells treated with 1 mM DTT and 1 mM bathocuproinedisulfonic acid (BCS) and harvested by centrifugaton. Pellets were washed twice with 150 mM NaCl. Cells were mineralized with 15.6 M HN03 (trace metal grade) for 1 h at 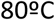 and overnight at 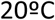. Digestions were completed by adding 2 M H_2_O_2_. Copper levels were measured by atomic absorption spectroscopy (AAS). When required, samples were diluted with 300 mM HNO_3_ before analysis by AAS.

### Copper content in cell fractions

Subcellular fractions, namely periplasm, cytoplasm, and total membrane fraction, were obtained using cells grown to mid-log phase in antibiotic-free LB medium. Aliquots (1.8 ml) were collected at indicated time points after addition of 0.5 mM CuSO_4_, treated with 1 mM DTT and 1 mM BCS and harvested by centrifugation. Pellets were washed twice with 1 ml 50 mM Tris-HCl, pH 7.6 and were kept on ice for no longer than 1 h prior to cell fractionation. Fractionation was performed following Ize *et al*. (2014) with modifications. The periplasmic fractions were obtained by resuspending cells in 0.5 ml of 200 mM MgCl_2_, 50 mM Tris-HCl, pH 7.6 and incubation for 10 min at room temperature with gentle shaking. After 5 min incubation on ice, periplasmic fraction was collected by centrifugation (8000 g for 10 min at 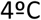). The resulting osmotically shocked cells and spheroplasts were washed with 1 ml 50 mM Tris-HCl, pH 7.6 and disrupted by sonication. Cells debris and unlysed cells were separated from cytoplasm and membrane fraction by centrifugation (2000 g for 15 min at 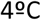). The procedure yielded >93±2 % cell lysis (n=5). Controls using anti-His-tag antibodies and cells expressing cytoplasmic His-tagged proteins could not detect the presence of cytoplasmic proteins in the periplasmic fraction (data not shown). Protein content was quantified in different fractions by Bradford (Bradford, 1976). Copper levels were measured by AAS as described above.

### Model development, simulations, and parameter estimation

All three models consist of six differential equations, the full models are provided in the Supplementary Text, and Supplementary Files in COPASI and SBML (Hucka *et al*., 2003) formats. The models were also deposited in the BioModels database (Chelliah *et al*., 2015) and assigned identifiers MODEL1804120001, MODEL1804120002, and MODEL1804120003. Computational analyses were carried out using the open source software COPASI (Hoops *et al*., 2006), version 4.21. The kinetic models are based on differential equations and solved using the LSODA integration algorithm (Petzold, 1983). Parameter values were estimated with COPASI’s parameter estimation task which minimizes the sum of squares of residuals (SSR) between model predictions and experimental data; minimization of the SSR was carried out with the simulated annealing algorithm (Corana *et al*., 1987) followed by the Hooke-Jeeves algorithm (Hooke and Jeeves, 1961). Parameter identifiability was carried out by profile likelihood analysis (Raue *et al*., 2009) using a previously described procedure (Schaber, 2012). All computations ran on desktop computers under Windows and Linux operating systems.

## Acknowledgements

This work was supported by a grant from the National Institutes of Health (R01GM114949 to J.M.A.). P.M. thanks the National Institutes of Health for funding work on COPASI (R01GM080219). The Authors declare no conflicts of interest.

## Author contributions

J.M.A, R.L. and P.M. were responsible for conception and design of the study, J.Q. and D.R. were responsible for acquisition of all experimental data. J.H.P. and P.M. carried out the computational work. All authors participated in data analysis, interpretation and writing of the manuscript.

